# Ribosomal intergenic spacers are filled with transposon remnants

**DOI:** 10.1101/2023.02.20.529178

**Authors:** Arnold J. Bendich, Scott O. Rogers

## Abstract

Eukaryotic ribosomal DNA (rDNA) comprises tandem units of highly-conserved coding genes separated by rapidly-evolving spacer DNA. The spacers of all 12 species examined were filled with short direct repeats (DRs) and multiple long tandem repeats (TRs), completing the rDNA maps that previously contained unannotated and inadequately studied sequences. The external transcribed spacers also were filled with DRs and some contained TRs. We infer that the spacers arose from transposon insertion, followed by their imprecise excision, leaving short DRs characteristic of transposon visitation. The spacers provided a favored location for transposon insertion because they occupy loci containing hundreds to thousands of gene repeats. The spacers’ primary cellular function may be to link one rRNA transcription unit to the next, whereas transposons flourish here because they have colonized the most frequently-used part of the genome.

**Author Summary:** The DNA loci containing the ribosomal RNA genes (the rDNA) in eukaryotes are puzzling. The sections encoding the rRNA are so highly conserved that they can be used to assess evolutionary relationships among diverse eukaryotes, yet the rDNA sequences between the rRNA genes (the intergenic spacer sequences; IGS) are among the most rapidly evolving in the genome, including varying within and between species and between individuals of a species, and within cells of an individual. Here we report the presence of large numbers of direct repeats (DRs) throughout the IGSs of a diverse set of organisms. Parasitic DNA and RNA elements often leave short DRs when they are excised resulting in “molecular scars” in the DNA. These “scars” are absent from the coding sections of the rDNA repeats, indicating that the IGSs have long been targets for integration of these parasitic elements that have been eliminated from the coding sections by selection. While these integration events are mostly detrimental to the organism, occasionally they have caused beneficial changes in eukaryotes, thus allowing both the parasites and the hosts to survive and co-evolve.

## Introduction

The IGS part of any rDNA locus is typical of rapidly-evolving satellite DNA (see below), variants of which are found elsewhere in the genome (near centromeres, e.g.), whereas the coding part of the very same satellite rDNA unit evolves extremely slowly [1–4]. Here we analyze rDNA sequences and propose a mechanism allowing such extreme differences in rates of evolutionary change in closely linked DNA segments. The coding part of rDNA is under heavy selection due to the vital function of ribosomes, whereas most of the IGS is the product of selfish DNAs that have colonized susceptible sections of the genome including the rDNA. Because rDNA loci contain hundreds to thousands of tandem copies, many of which are actively transcribed, they present large targets for integration of mobile genetic elements.

Maps of individual rDNA repeat units (Fig 1) include genes for the large, small, and 5.8S rRNA subunits flanked by two external transcribed spacers (3’ ETS and 5’ ETS) and two internal transcribed spacers (ITS1 and ITS2), as well as an IGS section that includes the 5S rRNA gene in some fungal species (*Saccharomyces cerevisiae*, *Flammulina velutipes*, e.g.), whereas in other fungi (*Schizosaccharomyces pombe*, *Yarrowia lipolytica*, e.g.), animals, plants, and protists the 5S genes are found outside of the rDNA loci, often on separate chromosomes [5–7]. What is most interesting from our perspective, however, is the IGS where, except for some short sequences representing transcription signals, several thousand bp of DNA are commonly depicted as lines without annotation, as if those sequences were so inconsequential as to be ignored. However, in recent work employing long-read sequencing, those sections were found to contain a hodge-podge of satellite DNA variants for which neither functional significance nor source was considered [6, 8]. In some algae, electron microscopy of nucleolar material showed transcriptionally-active tandem rDNA repeat units with long IGS sections, whereas other tandem units from the same nucleolus had no discernable IGS (Figs 2A−D; [9]). In some species IGS sections exceed 10 kb in length (*Homo sapiens*, e.g.), while in most species IGSs are only a few kb in length. Furthermore, many species have multiple variants of IGSs, with some longer and some shorter than the mean length of the rDNA population [3,4,10]. Such variability makes any functional contribution of the IGS to cellular phenotype difficult to discern.

**Fig 1.**
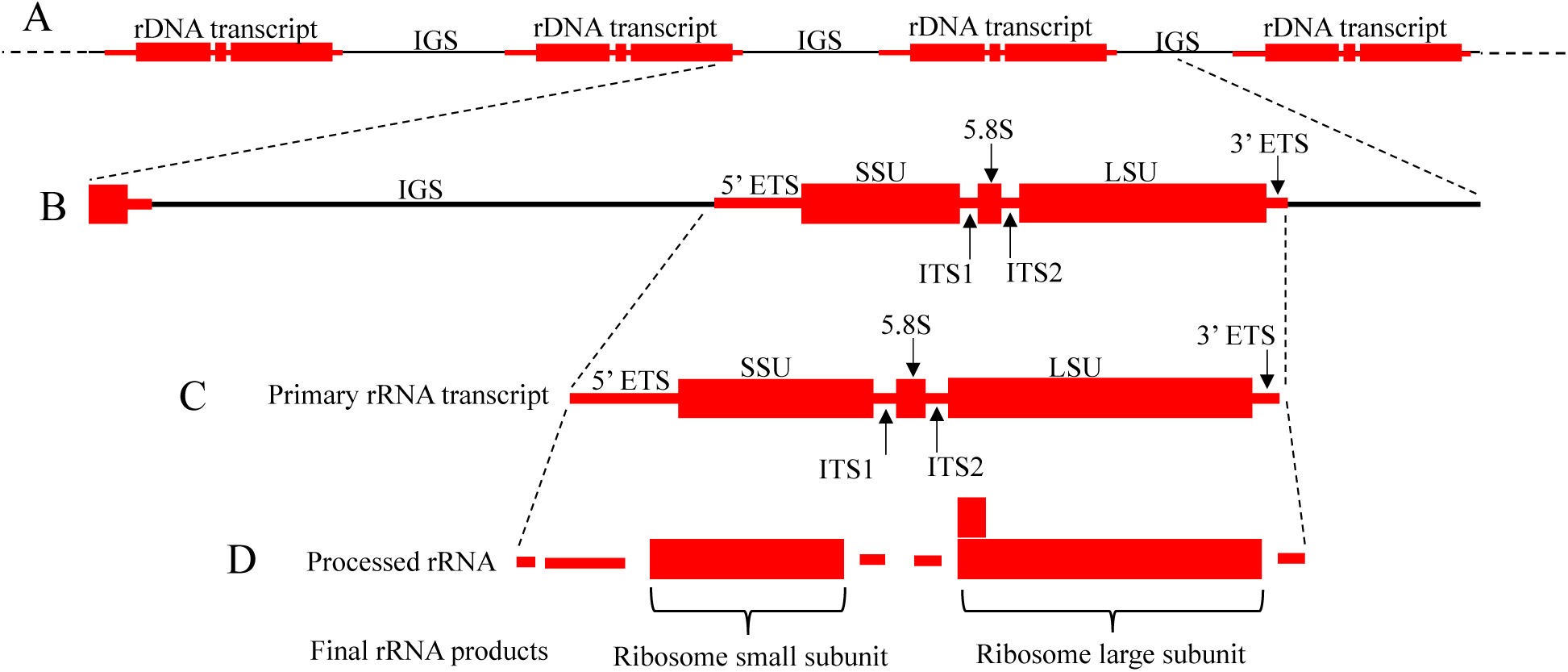
Ribosomal DNA locus. (A) Tandem array of eukaryotic nuclear rDNA showing four rDNA transcription units (red font) and IGS sections (black font). (B) A single rDNA unit with an IGS, a 5’ ETS, an SSU gene, two internal transcribed spacers (ITS1 and ITS2), a 5.8S gene, an LSU gene, and a 3’ ETS. Some eukaryotes also have a 5S rRNA gene within the IGS. (C) Primary rRNA transcript. (D) The rRNA after processing. All spacers are removed as the SSU, 5.8S, and LSU rRNAs are folded and assembled into the ribosome, with the addition of ribosomal (and other) proteins (not shown). [Note: In Bacteria and Archaea the rDNAs are dispersed in the genome, usually containing a 5’ ETS an SSU gene, an internal transcribed spacer (often containing one of more tRNA genes), an LSU gene, and a 3’ ETS. They have no IGS sections.

**Fig 2.**
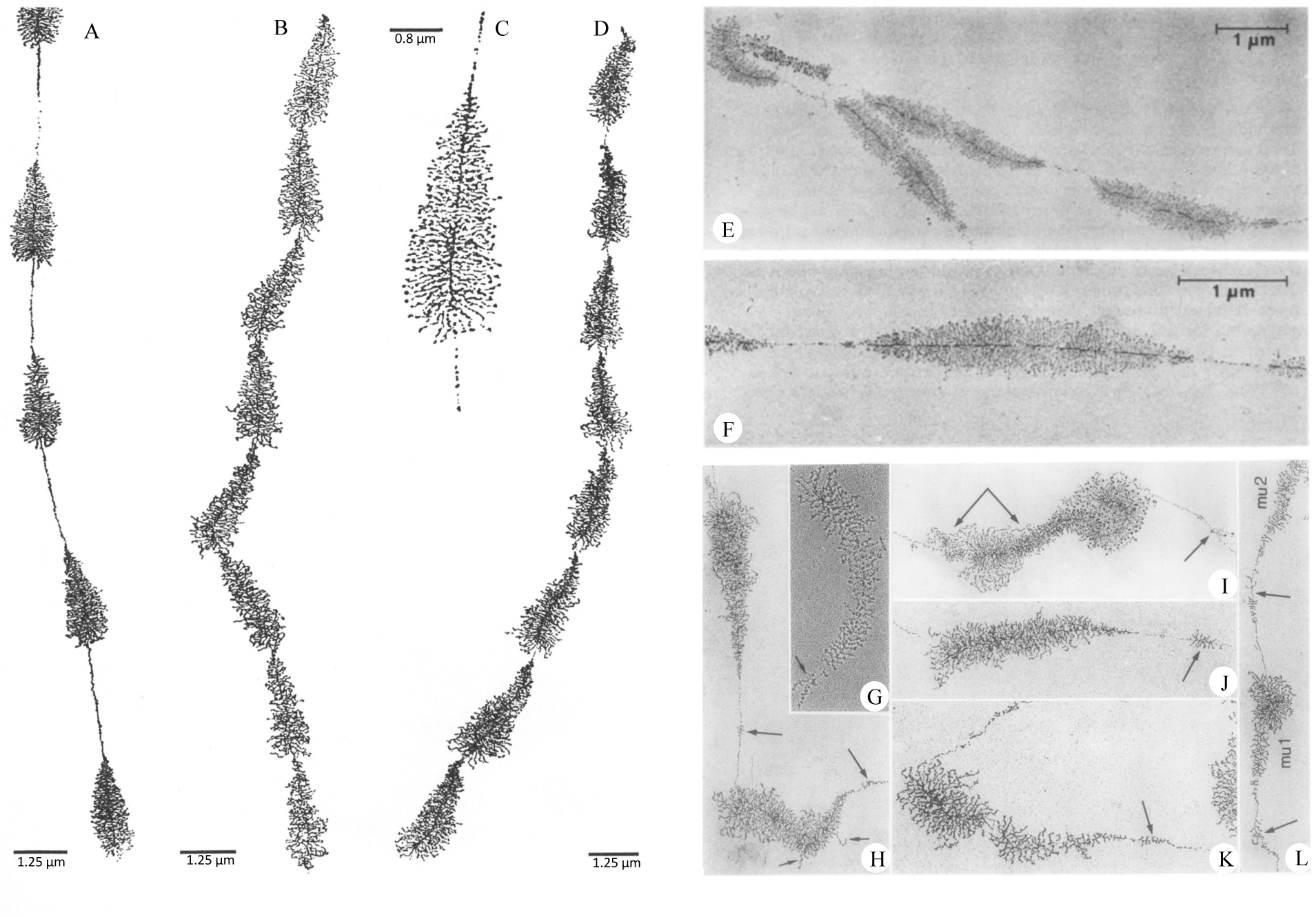
Transmission electron micrographs of rRNA transcription. (A-D) are from the green alga *Batophora oerstedii*, demonstrating rDNA repeats with long IGSs (A) and short IGSs (B and D). (A) and (B) are from the same individual nucleolus. A higher magnification is shown in (C). Micrographs (E) and (F) are from the green alga *Acetabularia exigua* and show gene repeat units that are inverted in head-to-head (or tail-to-tail) copies. Micrographs (G−L) show additional variations in the rDNA section. These include “tufts” (denoted by long single arrows) that indicate transcription within the IGS, as well as extensions of rRNA (G−L). H, K, and J are from the alpine newt *Triturus alpestris*; (G) and (L) are from the palmate newt *Triturus helveticus*; and (I) is from the house cricket *Acheta domesticus*; [18]). Micrographs are from: [9,18,19], with permission.

As will be described below, in prokaryotes the IGS appears to be absent and rDNA copy numbers are low. In eukaryotes IGSs are prominent and of variable length, and rDNA copy numbers can be high and vary greatly even among cells of an individual.

Our objective here was to reexamine rDNA sequence data so as to elucidate the structure and raison d’être of the IGS sequences that have previously been inadequately studied. Why are these seemingly paradoxical sequences and their divergent variants commonly found among diverse eukaryotes? Do these satellites contribute to cellular phenotype? What biochemical mechanisms can explain their genesis and rapid rates of change in sequence and copy number? How can we account for the strikingly different rDNA properties in eukaryotes and prokaryotes?

### Satellite DNA

When whole-cell DNA is analyzed by CsCl density gradient centrifugation, the major band is sometimes accompanied by secondary bands of either lower and/or higher density that typically differ in base composition from the major band: hence, “satellite DNA” (satDNA). In contemporary usage “satellite” no longer implies a particular base composition or the same sequence orientation between neighboring repeat units and may include “higher-order repeat units” containing subrepeats and extraneous sequences within the repeating unit [11]. SatDNA may contain tandemly-repeating sequences varying from as little as 2−6 base pairs (bp; termed microsatellites) to hundreds of bp to several thousand bp (macrosatellites), although these designations are rather arbitrary [12,13,14]. The degree of similarity in the tandem “repeats” varies among species, tissues, and even chromosomes of an individual cell. It is the tandem arrangement of repeating units—usually imperfectly repeating units—that best describes this type of DNA. Satellites are generally concentrated in sections of chromosomes near the centromere, telomeres, and in interstitial heterochromatic parts of chromosomes. Satellites could potentially destabilize chromosome structure if they were to participate in recombination, although this threat is usually suppressed.

Simple-sequence satDNAs can serve an important cellular function, such as in capping the ends of chromosomes in many eukaryotes. However, reverse transcriptase (a hallmark of retrotransposons) is also involved in this telomerase-dependent process. In *Drosophila*, tandem copies of retrotransposons can serve as telomeres [15]. Another example of simple-sequence usage is its contribution to the multi-protein kinetochore that connects chromosomes to kinetochore microtubules during chromosome segregation prior to cell division. This simple-sequence DNA is transcribed to a noncoding RNA needed for kinetochore assembly with other components, including DNA units in a cruciform structure, the nucleosomal histone H3 protein variant centromeric protein A (CENP-A) and other proteins, as proposed by Thakur et al. [14].

Tandemly-repeating simple-sequence DNA repeats can be created from parts of complex-sequence transposons. This conclusion applies to both animals and plants and to DNA transposons and retrotransposons, including LINEs and SINEs [12, 16]. The inference that such DNA can move around the genome is supported by several observations ([17], and references therein). Such DNA can be found in different sections of one or more chromosomes: High-copy tandem arrays are located in constitutive heterochromatin and outside of it in either low-copy arrays, single monomers or monomer fragments, and as short arrays within mobile DNA elements. The same or related unit sequence can be found both within high- and low-copy locations and as short arrays within MITE and Helitron transposons [12]. These examples show that DNA sequences originating as or generated by genomic parasites can later become indispensable to a host organism and illustrate the “bargain” struck between parasite and host.

### Historical Perspective

Prior to about 1990, the principal tools used to analyze rDNA were electron microscopy, restriction endonuclease digestion, and blot-hybridization. Three surprising conclusions were drawn from these early studies. First, the rRNA coding sections were highly conserved among eukaryotes, whereas the IGS sections evolved rapidly even among closely-related species [3, 4]. Second, the length of the IGS could vary among species, individuals in a population, and even during development of an individual plant or animal, whereas the rRNA coding section was not variable. Third, the rDNA copy number per genome was similarly variable, leading to the conclusion that there were more copies of rDNA than needed to support growth and development of the individual [3, 4].

After 1990, large numbers of rDNA sequences became available that confirmed the earlier generalizations. But the IGS sections were found to consist of subrepetitive sections and segments of unknown identity, so that it became difficult to map the coding and IGS sequences within the same rDNA repeat unit. This difficulty has only recently been overcome for a few species by using long-read sequencing methods.

Sections of DNA that are being transcribed have been visualized by annealing the nucleic acids to a thin film of nitrocellulose on an electron microscope grid, and then shadowing with palladium and/or platinum. These “spreads” are useful in observing the nuances of rDNA transcription and in characterizing IGSs. Individual nucleoli were found to be simultaneously transcribing rDNA with variable lengths of IGS (Figs 2A−D), including rDNA repeats that have IGSs that are less than 350 bp in length in addition to those between 6.5 and 7.0 kb [9]. Unusual transcriptionally-active rDNA repeats were also observed: head-to-head dimers; truncated rDNA units (Figs 2E and F; [18, 19]); and short transcripts (i.e., tufts) within the IGS sections (long arrows in Figs 2H−L).

Blot-hybridization, electron microscopy, and sequencing studies of the IGS reported long tandem repeats (TRs) within most species that often varied in number within a species and within individuals. The IGS in *Vicia faba* contained 0−23 (or more) 325-bp TRs and variable numbers of 150-bp TRs [3,4,20,21]. Seven other species of *Vicia* also exhibited variation in IGS length [4]. The IGSs within *Pisum sativum* had 0−30 (or more) 180-bp TRs [9,22,23,24]. Furthermore, in *P. sativum*, there were two rDNA loci, one with only two IGS size classes and the other with a wide range of IGS size classes [24]. The IGS in *Arabidopsis thaliana* also exhibited variability in repetitive elements [25]. R-looping studies of *V. faba* rDNA hybridized to rRNAs showed that the length of one IGS was unrelated to the lengths of the adjacent IGSs, exhibiting a seemingly random organization of IGS size lengths along the chromosome [4]. In the same study, individual plants exhibited a 95-fold variation in rDNA copy number, from 230 to 21,900 copies per haploid genome. Within an individual, a 12-fold variation in copy number was measured, and large copy number variations were reported from one generation to the next. Variation in copy number has been reported in many species (including humans).

In summary, early research on the IGS revealed great variability in amount, spacing, and sequence organization, so that its cellular function, if any, was perplexing. By contrast, its flanking coding sections were highly conserved as expected, considering their translational importance. Here we report that the extreme variation in the IGS sections was the result of frequent insertions and deletions of transposons.

## Results and Discussion

### Tandem Repeats

TRs were present in the IGS of all species examined, ranging in size from about 10 to more than 2000 bp (Figs 3A and B; Table 1). Each of the TR sections was flanked by a pair of short direct repeats (DRs; 3−8 bp each), suggesting that they originated from a transposition event. Some individual repeats within the TRs had the same DR sequences at each of their flanks (Fig 3: R in *O. sativa*; R2 and R3 in *V. faba*; R1 in *A. thaliana*; R1 and R2 in *D. funebris*; and R1 in *G. gallus*). TRs were also found within the 5’ ETS in three of the plants examined (*A. thaliana*, *V. faba*, and *V. sativa*) and one animal (*C. nozakii*). Two species had a single TR type (*O. sativa* and *P. sativum*). Four had two types of TRs (*A. thaliana*, *F. kerguelensis*, *C. nozakii*, *D. funebris*), four had three types (*V. sativa*, *F. velutipes*, *G. ultimum*, *G. gallus*), one had five types (*V. faba*), and *H. sapiens* contained two major TRs and dozens of Alu/SINE elements of various lengths. In the basidiomycete, *F. velutipes*, a 5S rRNA gene was also present in the IGS and it was flanked by DRs, dividing the IGS into two sections (IGS1 and IGS2). The maps in Figure 3 represent only one version of each IGS and ETS section. As mentioned above, length variants are known for most of the species. Individual organisms, tissues, and loci may contain all or most of the IGS size variants, or only a limited number. Each of the individual repeats within each TR section was nearly identical, except some repeats in *C. nozakii*, *O. sativa*, and *A. thaliana* were truncated (Figs 3A and B; Table 1), and most of the Alu/SINEs in the human IGS were truncated variants (Fig 3B). Most of the TRs were unique among the species and within an IGS, although there were some similarities among closely-related species (e.g., *V. faba* and *V. sativa*). Primate IGS sequences exhibit sections that are somewhat conserved among all species and genera, while still containing long sections unique to each [26]. Overall, sections of the IGS and ETS exhibit vertical inheritance within a species or genus, but sequence similarity declines rapidly above the genus level.

**Fig 3.**
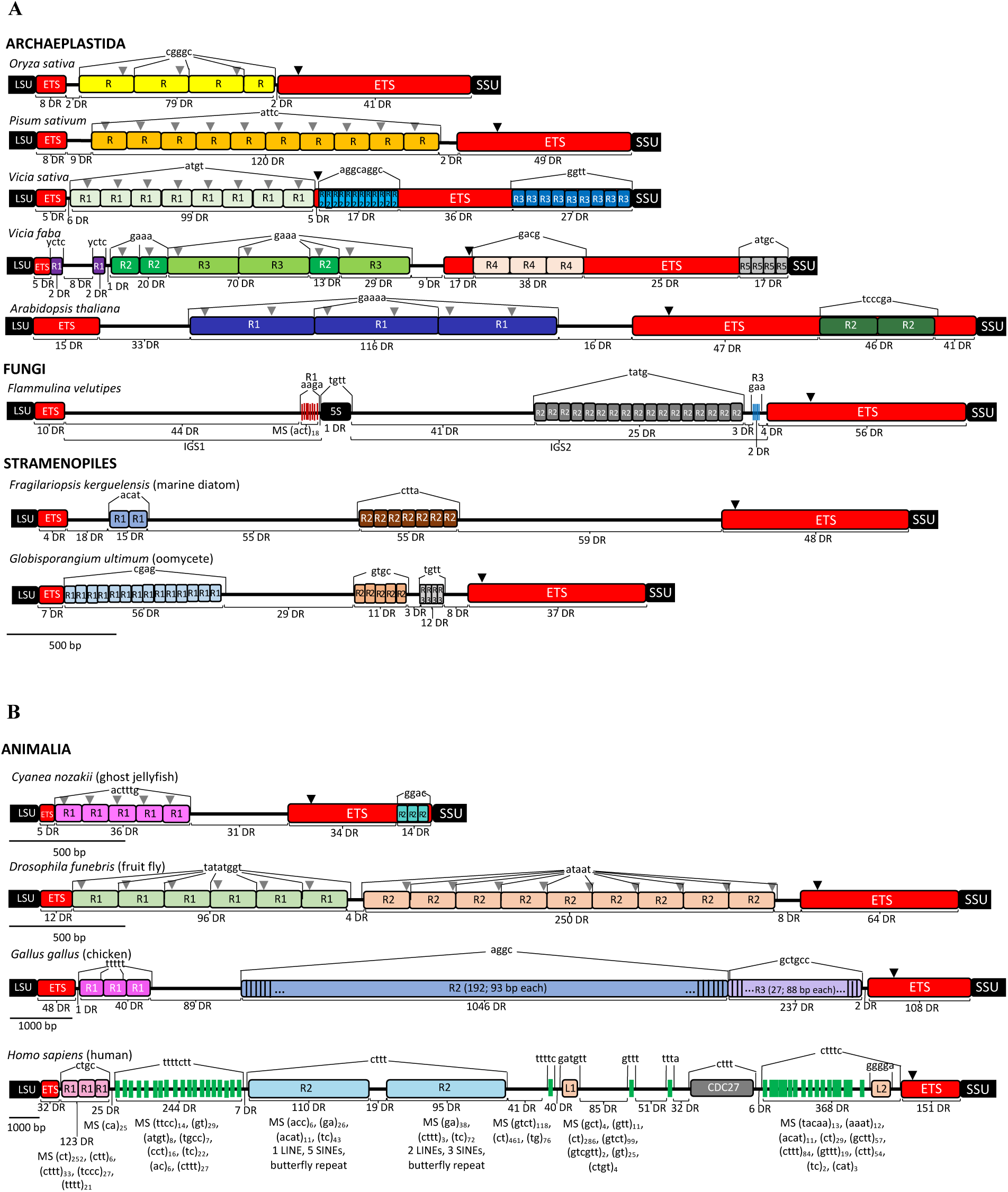
Maps of nuclear rDNA IGS plus 3’ ETS (red rectangles on left) and 5’ ETS (red rectangles on right) segments. (A) Archaeplastida, Fungi, and Stramenopiles. (B) Animalia. Sections with TRs ≥20 bp are shown by colored rectangles or colored lines. Each of these sections is flanked by short direct repeats (DRs) indicated above each repeat section, but some of the same DRs also are found at the borders of each of the individual TRs within those TR sections (*O. sativa* R repeats, *V. faba* R2 and R3 repeats, *A. thaliana* R1 repeats, *D. funebris* R1 and R2 repeats, and *G. gallus* R1 repeats). In *A. thaliana*, for example, the group of three R1 repeats is flanked by gaaaa, as is each individual R1 repeat, whereas the group of two R2 repeats is flanked by tcccga. The numbers of short DRs (2−10 bp each) are indicated below each of the sections, including within the TRs and the ETSs. The black triangles within the 5’ ETS are the locations of promotor sequences that also have upstream RNA polymerase binding sites approximately 30 bp upstream from the promoter. The gray triangles in some TRs are locations of sequences nearly identical to the promoter that also have upstream binding sites. Thus, they are similar to promoters for RNA pol I (including a “TATA” sequence followed by several G residues and a CGCC upstream binding site). Promoters for RNA pol II or pol III were absent. Alu/SINEs in the human IGS and ETS segments are shown. Microsatellites also are shown for *H. sapiens* because of their lengths, total numbers, and sequence diversity. Large numbers of microsatellites are also present in chicken, but are much less numerous in all the other species.

**Table 1.** IGS and ETS repeat characteristics.

| Taxon | Species | Genome size (Gb) | IGS+ETS length (kb)<sup>a</sup> | Number of DRs | DRs per kb of the IGS+ETS | Percent segment length as repeats<sup>b</sup> |  |  |  | TR type | TR lengths (bp) | Mean length (bp) |
| --- | --- | --- | --- | --- | --- | --- | --- | --- | --- | --- | --- | --- |
|  |  |  |  |  |  | 3' ETS | IGS | 5' ETS | TOTAL |  |  |  |
| Archaeplastida | *A. thaliana* | 0.14 | 4.7 | 316 | 67 | 42.2 | 52.5 | 59.7 | 55.4 | R1 | 616-621 (547)<sup>c</sup> | 618 |
|  |  |  |  |  |  |  |  |  |  | R2 | 304-310 | 307 |
|  | *O. sativa* | 0.40 | 2.0 | 133 | 66 | 58.0 | 49.1 | 47.3 | 46.9 | R | 255-265 (115)<sup>c</sup> | 259 |
|  | *P. sativum* | 4.5 | 2.75 | 189 | 68 | 47.9 | 50.3 | 44.5 | 48.0 | R | 139-212 | 175 |
|  | *V. faba* | 13.0 | 3.25 | 192 | 57 | 43.5 | 59.3 | 48.5 | 53.7 | R1 | 28 | 28 |
|  |  |  |  |  |  |  |  |  |  | R2 | 144-145 | 144 |
|  |  |  |  |  |  |  |  |  |  | R3 | 319-328 | 324 |
|  |  |  |  |  |  |  |  |  |  | R4 | 163-166 | 165 |
|  |  |  |  |  |  |  |  |  |  | R5 | 59-71 | 65 |
|  | *V. sativa* | 1.8 | 2.75 | 146 | 52 | 46.4 | 59.2 | 43.5 | 49.9 | R1 | 127-207 | 166 |
|  |  |  |  |  |  |  |  |  |  | R2 | 20-21 | 20 |
|  |  |  |  |  |  |  |  |  |  | R3 | 62-67 | 66 |
| Fungi | *Fl. velutipes* | 0.36 | 4.1 | 229 | 55 | 47.7 | 50.9 | 44.4 | 49.0 | R1 | 12-17 | 14 |
|  |  |  |  |  |  |  |  |  |  | R2 | 37-50 | 47 |
|  |  |  |  |  |  |  |  |  |  | R3 | 7-9 | 8 |
| Stramenopiles | *Fr. kerguelensis* | 0.30 | 4.0 | 211 | 52 | 53.5 | 54.9 | 55.6 | 54.9 | R1 | 81-83 | 82 |
|  |  |  |  |  |  |  |  |  |  | R2 | 65-71 | 68 |
|  | *G. ultimum* | 0.42 | 2.8 | 166 | 58 | 68.2 | 50.5 | 49.2 | 50.8 | R1 | 59-75 | 69 |
|  |  |  |  |  |  |  |  |  |  | R2 | 38-47 | 43 |
|  |  |  |  |  |  |  |  |  |  | R3 | 18-24 | 21 |
| Animalia | *C. nozakii* | 0.15 | 1.8 | 122 | 67 | 67.7 | 43.7 | 49.5 | 46.6 | R1 | 116 (87)<sup>c</sup> | 116 |
|  |  |  |  |  |  |  |  |  |  | R2 | 61-75 (37)<sup>c</sup> | 68 |
|  | *D. funebris* | 0.20 | 4.0 | 252 | 63 | 40.7 | 58.2 | 60.2 | 57.5 | R1 | 238-240 | 239 |
|  |  |  |  |  |  |  |  |  |  | R2 | 227-237 | 232 |
|  | *G. gallus* | 1.2 | 15.2 | 1544 | 102 | 59.4 | 68.2 | 61.2 | 67.4 | R1 | 345-448 | 395 |
|  |  |  |  |  |  |  |  |  |  | R2 | 92-94 | 93 |
|  |  |  |  |  |  |  |  |  |  | R3 | 78-91 | 88 |
|  | *H. sapiens* | 3.1 | 31.5 | 1429 | 45 | 59.3 | 62.7 | 56.3 | 62.4 | R1 | 715-800 | 758 |
|  |  |  |  |  |  |  |  |  |  | Alu/SINEs | 18-242<sup>d</sup> | 89 |
|  |  |  |  |  |  |  |  |  |  | R2 | 2008-2056 | 2032 |

<sup>a</sup>As indicated in the main text, these lengths often vary within and among individuals.
<sup>b</sup>Repeats include DRs and microsatellites.
<sup>c</sup>Numbers in parentheses indicate truncated TRs.
<sup>d</sup>Length range represents full-length and truncated Alu/SINEs.

### Short Direct Repeats and Microsatellites

Large numbers of DRs (dozens per kb: Table 1; Figs 3A and B) were found throughout the IGS and ETS sections in all species. DRs ranged in length from 2 to 10 bp (per monomer), most of which were in pairs, and occasionally three or more DRs occurred within a span of 4−40 bp. In most cases, the individual repeats comprising the DRs were adjacent to one another, although many were separated by several base pairs. Some DRs overlapped other unrelated DRs, possible indications of insertion partially within an existing unrelated insertion element. This was frequently observed in the *G. gallus* IGS. The number of DRs/kb in the IGSs + ETSs ranged from 45 (*H. sapiens*) to 102 (*G. gallus*), with a mean of 63 DRs/kb. The total number of DRs in the IGSs + ETSs ranged from 133 (*O. sativa*) to 1544 (*G. gallus*), with a mean of 409 DRs (193, excluding *G. gallus* and *H. sapiens*).

The abundance of microsatellites ranged from a few in most species to >1000 in the vertebrates. In *O. sativa* a “gc” monomer was repeated 3−4 times, and in *V. faba* a “cg” monomer was repeated 5 times. *Arabidopsis* had “gt” up to 4 times, “cg” up to 6 times, and long tracts of “a”. The fungus and stramenopiles also had a few simple-sequence repeats. The jellyfish had “gt” repeated 4−5 times, while the fruit fly had “at” dinucleotides repeated up to 7 times and “tg” up to 3 times. The number and lengths of microsatellites in the chicken IGS were much greater than those in plants, fungi, and stramenopiles: at least 10 different microsatellite types (e.g., “cg”, “ccgg”, “ga”) repeated up to 8 times each at many sites and comprising hundreds of nucleotides, as well as long tracts of repeating t, c, and g monomers. In the human IGS, microsatellite expansion was even greater. At least 40 different microsatellite types were found (Fig 3B), some repeated hundreds of times representing thousands of nucleotides. The IGSs in chicken and human are much longer than in the other 10 species, with DRs and microsatellites accounting for this difference.

Although the numerous DRs described above were identified by manual inspection of sequences separated by no more than 40 bp, larger numbers of DRs were found when a bioinformatic approach was used (see Materials and Methods). A program designed to find DRs (2-10 bp per monomer) without the constraint to be separated by a short distance resulted in locating many more DRs in the IGS and ETSs (Table S1, and Figs S1-S3). When compared to random sequences of the same G+C percentage, the IGS+ETS sections in all species had significantly more DRs than the random sequences at the p<0.01 level (Table S1), with the following exceptions: 2-4 bp repeats in *O. sativa* and 2 bp repeats in *C. nozakii*. By contrast, the numbers of DRs within the rDNA coding sequences were not significantly different from random sequences.

### Summary of IGS data

For each of the 12 species investigated, our analysis revealed that the IGS and ETS sections were composed mainly of TRs and DRs that may have originated from the entry and exit of transposons. For each species all parts of the IGSs conformed to this pattern, so that no species required an unannotated section to complete the map.

### Parasitic Sequences

There are two previously-described types of parasitic DNAs that can proliferate within the nuclear genome. The first utilizes a transposase encoded by an autonomous parasite to mobilize itself as well as truncated nonautonomous versions of itself (e.g., LINEs and SINEs, respectively). The second type, exemplified by MITEs, utilizes the transposase from other transposons (not classified as MITEs) for their own proliferation: a parasite of a parasite. For both types, most of the enzymes required for transposition and proliferation are encoded by host cell DNA and used by the parasitic DNA. SatDNA found in the IGS may represent a third type in which there is no sequence-specific transposase and all the enzymes required for proliferation are encoded by the host cell. The connection between transposons and satDNA has been reported for numerous taxonomic groups of animals and plants [12,15,27]. A transposon transcript inserts at a break in the DNA, followed by invasion and reverse transcription, which is an error-prone process (Fig 4; [28]). A host DNA polymerase then synthesizes the opposite strand. Target site duplications (TSDs), which are DRs, are formed on both flanks of the insert during synthesis and integration.

**Fig 4.**
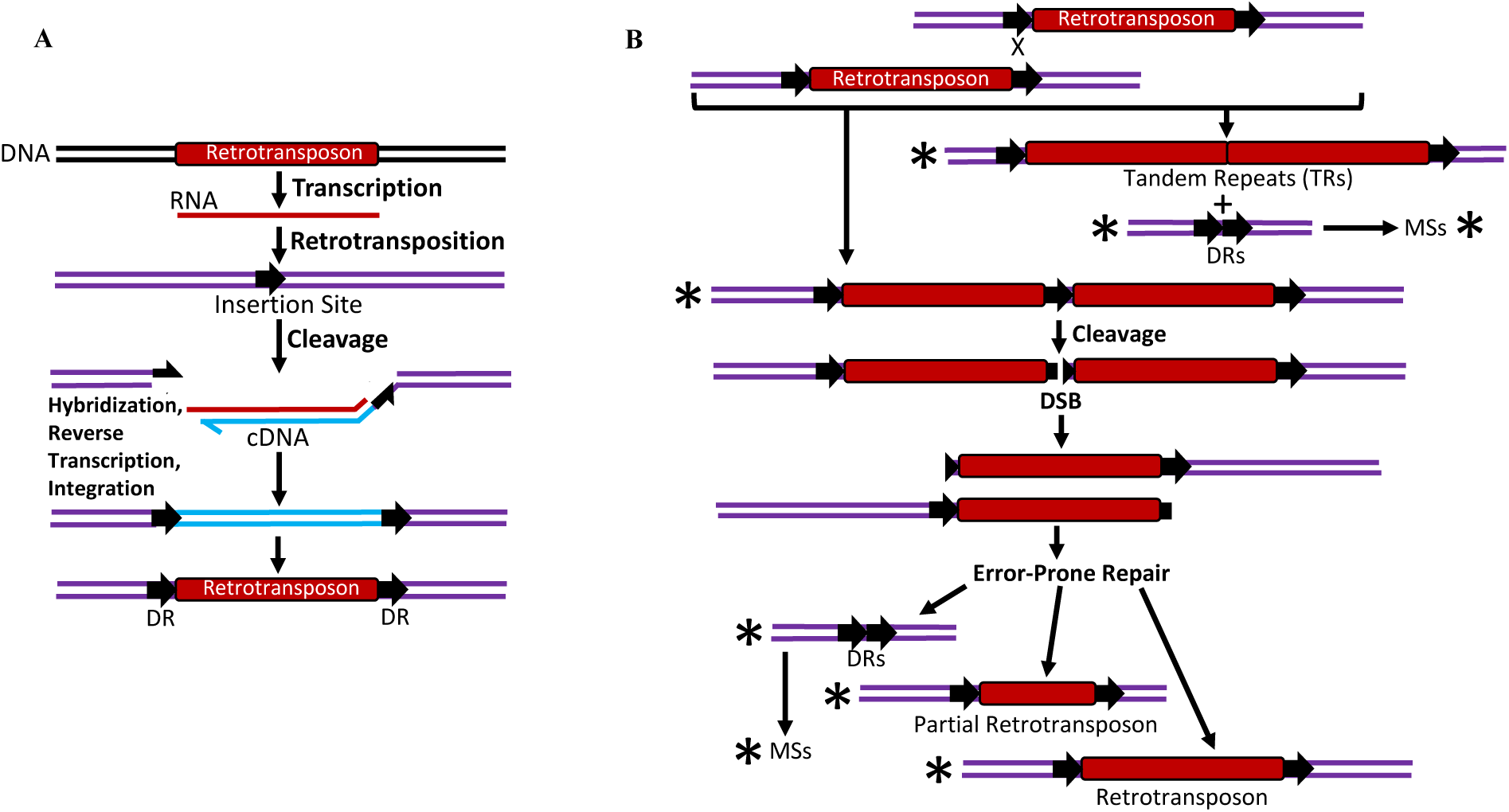
Proposed mechanism of insertion, duplication, and elimination of retrotransposons in the rDNA IGSs and ETSs. (A) When retrotransposons move into a site, they do so at DNA breaks they create or at preexisting breaks. Retrotransposons insert as single-stranded RNA, followed by reverse transcription, and then host DNA polymerase produces the complementary strand to complete integration of a DNA copy of the retrotransposon. During transposition, a TSD (a DR, i.e.) is created that flanks the retrotransposon. (B) Recombination is active in the rDNA loci. This can cause tandem duplications of the inserted sequences, with or without retention of the DRs, and can also lead to elimination of part or all of the retrotransposon. The duplications have resulted in the TR blocks, still flanked by the original DRs. Elimination of the transposons leads to a section of the IGS that lacks all or most portions of the insert but leaves the remnant DRs. Both results are documented among the sequences used in this study (structures with asterisks). The insertion process also may create a double-strand DNA break (DSB) that must be repaired. The repair process may generate single-stranded DNA sections vulnerable to damage, use error-prone DNA polymerases, and involve DNA polymerase slippage and template switching [28], leading to the formation of tandem repeats, partial inserts, elimination of the inserts, isolated DRs, and microsatellites (MSs) [12, 15]. All of these have been found (indicated by asterisks) in the IGSs of the species examined. This scheme is based on the work of Deininger [29].

The DRs that we identified within the IGSs represent the remnant TSDs from ancient transposition events. When the transposons are eliminated from the site via recombination or other mechanisms (Fig 4B; [29]), they can leave behind telltale signs of their visitation, including DRs. While the human IGS contains mainly Alu/SINE elements, the type of transposons that have been found in other species’ IGS and ETS sections is unclear. However, DNA encoding lncRNAs, miRNAs, and sRNAs have been shown to transpose within genomes [12], and the R1 repeats in the *A. thaliana* IGS have sequence similarities to some of these (Fig 3A). The 5S genes in most species reside in loci separate from the large rDNA locus, and many species have multiple 5S gene loci. However, 5S genes have been found in the IGS of many fungal species (e.g. *F. velutipes*; Fig 3A). We identified DRs flanking the 5S gene in the *F. velutipes* IGS, indicating that transposition of these genes is a likely cause for the different locations.

The large number and variety of repetitive sequences found in the IGS and ETS indicates that transposition events have occurred often, probably over billions of years [30]. They appear to have had minimal functional effects on these sections. The strongest evidence that the absence of an IGS has no effect on rRNA production comes from transcriptional activity revealed in Miller spreads (Figs 2A−D) and blot-hybridizations that demonstrate extremely short IGSs. These data indicate that the entire IGS (except for the proximal promoter) can be removed from the rDNA, thereby reducing the target for transposon invasion. For most species, however, the host cell apparently tolerates the transposons, their remnants, and their elongated IGSs.

### Copy number of rDNA

Eukaryotic cells contain many thousands to millions of ribosomes and many copies of rDNA. For example, in *D. melanogaster*, *H. sapiens*, *S. cerevisiae*, and *V. faba*, typical rDNA copy numbers range from 140 to 250 per haploid genome, although individuals can survive with fewer than half of those copies [3,4,31,32,33]. Copy number variation within a population of phenotypically indistinguishable individuals may exceed these mean values by 10-or even 100-fold [4]. It seems unlikely that these excessively-large numbers are useful to the host cell.

On the other hand, the repeating rDNA unit comprises not only the coding sequences, but the intergenic section dominated by transposons and their variants. The main beneficiary of “excess” rDNA copies may thus be the transposons enlarging their numbers and their target sites. The transition from scattered rDNA units in prokaryotes to tandem units in eukaryotes may well have been initiated by the insertion of repetitive transposons and DRs at the flanks of the rRNA genes.

### IGS Function

The promoter section for the rRNA genes is located within the 5’ ETS (black triangles in Fig 3, e.g.). Although similar sequences exist in some of the TRs (gray triangles in Fig 3), their function has not been investigated in any of these 12 species. It is therefore unclear how, or indeed whether the TRs are useful in producing the rRNAs in ribosomes or whether they are simply products of transposition and recombination. The length differences among IGS and ETS sections vary by as much as 15-fold (*H. sapiens* versus *O. sativa*) and stunningly from 5-fold to 25-fold within a species (*V. faba* and *P. sativum,* respectively). The smallest IGS sections are devoid of any of the largest TRs that contain putative promoters [3,4,10], suggesting their lack of cellular function in expression of the adjacent genes. Additionally, electron microscopy demonstrated the transcription of rDNA repeat units that were separated by long as well as very short IGS sections in the same nucleolus (Figs 2A−D; [9]).

In *A. thaliana*, however, sequences in the IGS TRs also align with sRNAs, lncRNAs, and miRNAs. These classes of RNAs are involved in regulating gene expression and have been shown to transpose to various genomic locations [27]. The tufts observed in electron micrographs (Figs 2G−L) of IGS sections might represent the production of RNAs affecting gene expression. Alternatively, these transcripts might represent the start of retrotransposition. Among some IGS sections in *A. thaliana*, gypsy-like LTR-retrotransposons and other retrotransposon sequences have been reported (NCBI accession number AC006837). Similarly, Alu/SINE, LINE, and LTR retrotransposons have been found in the human IGS, suggesting that at least some portions of IGSs originated from retrotransposons. Whether these IGS sequences modulate gene expression in *A. thaliana* or other species is unknown, although their movement into the IGS is apparent. Both the large number of DRs and the within-species variability of IGS length show that a mutually-tolerable interaction between host and parasite has a lengthy history among eukaryotes.

When analyzed statistically, compared to random sequences of the same base composition, the IGSs all had significantly more DRs than the random sequences (see Supporting Information Table S1, and Figs S1-S3). Somewhat surprising was the finding that the 3’ and 5’ ETSs had significantly more DRs than the random sequences. Both sections are transcribed but are processed out of the mature rRNAs (Fig 1). The 3’ ETS contains sequences responsible for transcription termination signals and the 5’ ETS has sequences responsible for the start of transcription. Most of the ETSs were adjacent to TRs, and the 5’ ETSs in four species (*Vicia sativa*, *V. faba*, *Arabidopsis thaliana*, and *Cyanea nozakii*) contained TRs, resulting in a wide range of lengths in the 5’ ETSs. Therefore, parts of the primary transcript represent transposon DNA inserted during evolution. When analyzed in the same way, the SSU and LSU sections contained no more DRs than were predicted from the random sequences. The SSU and LSU may also have experienced transposon insertions, but heavy selection to maintain functional ribosomes has caused the extinction of the rDNA repeats and/or organisms with appreciable numbers of the mutant rDNAs.

### Benefits and Beneficiaries of the IGS

The IGSs of some species contains sequences that are clearly beneficial to that species, although such sequences account for only a small part of the IGS that is dominated by transposon fragments and rapidly-evolving repeats (Fig 3). We now address the question of how such a mishmash of seemingly useless DNA may have originated. An IGS may contain sequences of three types: sequences that benefit (1) themselves as selfish DNA; (2) the cell; and (3) both themselves and the cell. In addition, the tandem rRNA coding units may not be separated by discernible IGSs, as in *Batophora* (Fig 2). Type 2 has been intensively studied in some animals and yeasts, and subrepeats of sequences affecting the transcription and maintenance of downstream rRNA-coding DNA have been identified [5,32,34]. For example, in cultured human cells IGS sequences transcribed in the antisense direction by RNA polymerase II can defend the cell during imposed stressful conditions [35]. The IGS in wild populations of *Tigriopus* copepods is exceptionally short (2.8 kb) and does not contain the subrepeat structure common to other eukaryotes, such as *Drosophila funebris* (Fig 3; [36]), so that possible IGS-mediated defense of rDNA would involve some other mechanism. The rRNA copy number ranged from 230 to 21,900 per haploid genome among 434 individual *Vicia faba* seedlings [4]. The individual with 230 may or may not carry an IGS that defends the cell during stress. Yet even if all 21,900 copies in the other individual encode functional (though unused) rRNA, this enormous rDNA copy number would likely be detrimental to the cell (see below) but increase copies of their parasitic sequences: type 1 IGS sequences.

When a pathogen sweeps through a population, not all individuals succumb to the infection. In prokaryotes, only ∼2% of the genome is comprised of mobile genetic elements and defenses against these invaders [37–39], whereas the eukaryotic genome is comprised mostly of repeated sequences [40]. Conceptually, the genes repairing transposon-induced damage in prokaryotes are strong alleles, whereas those in eukaryotes are weak alleles that allow the parasitic sequences to proliferate. As described for satDNA, a host that survived an rDNA insertion may later evolve a modified version for its own benefit: a type 3 IGS. This derived benefit evidently balances the burden of repairing the additional DNA damage and chromosome-disruptive recombination attending “extra” rDNA copies [34] sporadically distributed among plant and animal species.

### The source of the IGS repeats

Prokaryotic genomes typically contain several rRNA operons, but these copies are not spaced by repeat-containing satellite DNA sequences as found in the IGS of eukaryotes. The history of the IGS may therefore be elucidated by considering the transition from prokaryotes to eukaryotes. The *E. coli* genome contains seven *rrn* operons, and the consequences of altering of this number show that: (i) All seven are required for rapid adaptation to changing environmental conditions; (ii) Too few copies cause R-loop formation, chromosomal breakage, and cell death; and (iii) Additional copies lead to increased recombination and deleterious chromosomal rearrangements [41, 42]. Thus, in its natural habitat, preservation of chromosomal integrity determines the optimal copy number of rDNA for this bacterium and, we assume, the same would hold for eukaryotes unless some feature of eukaryotic life drives the rDNA copy number beyond that optimal for perpetuation of the organism. In our opinion, the chromosomal damage and instability created by transposons is strongly suppressed in prokaryotes but weakly suppressed in eukaryotes, leading to the repeats that dominate the IGS.

### Consequences of the IGS repeats

The rDNA is thought to be the most unstable genic part of the eukaryotic genome, and this copy number instability may benefit the organism in times of stress and during development, although copy number instability may also lead to medical disorders in humans [32]. When tandem 325-bp repeats from the IGS of *Vicia faba* (R3 in Fig 3) were introduced into *E. coli*, recombination occurred frequently among the repeat units [43], suggesting that the IGS is a recombination “hot spot” that may cause copy number instability in eukaryotes. Thus in eukaryotes, but not prokaryotes, copy number instability driven by the IGS may benefit or harm the organism. In either case, the parasitic sequences in the IGS proliferate and spread within the genome.

### Generating repeats in the IGS and possibly elsewhere

The insertion of a linear transposon (e.g., a retrotransposon), creates a double-strand DNA break (DSB) at the target site. We suppose that repair of this DSB resembles the repair of one-ended DSBs by break-induced repair (BIR) and the related synthesis-dependent strand annealing process in yeast [44, 45]. BIR involves persistent exposure of ssDNA, secondary (non-B-form) DNA structures, inverted repeat-induced polymerase slippage, error-prone DNA synthesis, short insertions/deletions, and mutagenesis [46]. Simple-sequence satellite DNAs with a local replication advantage may thus expand within the IGS (Fig 4), spread to other genomic locations by recombination, and act as selfish mobile elements without a dedicated transposase found in classical autonomous transposons.

Could this mechanism for generating the IGS repeats also apply to the rest of the genome? To address this question, we need to consider how the data were analyzed. The numerous DRs of length 2−10 nt depicted in Fig 3 were identified by visual inspection of thousands of bp of IGS sequences available in data bases. A repeat-search algorithm also was used to identify these short IGS repeats. This process identified many more DRs because the distances between the DRs was not considered. However, it was ineffective at searching sections of more than about 30 kb, so it would not be appropriate for genome-wide searches. Until appropriate algorithms are available to search entire genomes [47], we cannot answer the question of the extent of DRs in genomes. There is, however, a possible mechanism by which enormous numbers of tandem repeats (satDNA) found in heterochromatic sections of chromosomes might be produced. RAD52 is a protein involved in BIR in the nucleus. In human cells defective for RAD52, BIR was found to re-replicate an affected segment of the genome [48]. The product of such re-replication is tandem copies in potentially great numbers found in centromeres, sub-telomeres, or any part of a genome.

When the total lengths of repeats (DRs + microsatellites) was compared to the total lengths of the IGS + ETS sections, repeats collectively account for 47–67% of the IGS + ETS (“TOTAL” column in Table 1), a range similar to that reported for the fraction of many plant and animal genomes attributed to repetitive DNA sequences [40, 47]. Such estimates, however, depend on arbitrary criteria for the definition of a “repeated” DNA sequence. For four land plants, the fraction of the genome classified as repeated sequences increased from about 10% to 55% as the temperature decreased in DNA hybridization kinetics assays, relaxing the criterion required for sequence repetition [49]. In our present analysis, perfect sequence identity was required to classify short sequences as “repeated”, so that the 47−67% values in Table 1 would increase if the criterion for repetitiveness were relaxed. Thus, in our opinion, the rDNA represents a microcosm of the rest of the nuclear genome.

### Future Direction

Several questions remain to be addressed regarding the evolution of the rDNA loci: Why is the density of DRs in the IGS+ETS so similar among eukaryotes?; Do the properties of the IGS and ETS described here extend to other diverse eukaryotes?; Can an algorithm be created to search for DRs at the genome level?; Can the power of yeast genetics be used to elucidate the origin and raison d’être of the IGS+ETS sequences?; Are there species of Bacteria and/or Archaea with eukaryotic-like rDNA repeats, that may indicate transposon-like activities?; Do the rDNA transcribed spacers in Bacteria and Archaea also contain DRs indicative of visitation by transposons?

### Concluding remarks

McClintock recognized two kinds of “shock” that genomes may experience [50]. For the first, preprogrammed responses are mobilized to protect the structural integrity of the genome, such as the heat shock response in eukaryotes and the SOS response in bacteria. For the second, unanticipated challenges are met in an unforeseen manner. We are concerned with the second of these, when a retrotransposon integrates at a site within the IGS and ETS parts of nuclear rDNA and creates a double-strand DNA break. In the ensuing havoc, sequences are altered and the break is repaired using some of the components that protect against the first type of genome shock. This defensive action may succeed, but the host cell incurs a metabolic burden by adding somewhat deleterious DNA to the IGS and ETS in the form of transposon and simple-sequence DNA. McClintock [50] wrote that “it is necessary to subject the genome repeatedly to the same challenge in order to observe and appreciate the nature of the changes it induces”, a statement of astonishing prescience that provides a simple explanation for the heretofore bewildering nature of the IGS. Whereas McClintock’s evidence came from color changes in the seed, the rDNA sequence evidence comes from the most frequently needed part of the genome—a remarkable realization.

## Materials and Methods

### Sequences Used

All nuclear rDNA sequences were retrieved from NCBI during early 2022. The 12 species examined were: Animalia [*Cyanea nozakii* (MH813455), *Drosophila funebris* (L17048), *Gallus gallus* (MG967540), and *Homo sapiens* (MF164258), [Archaeplastida [*Arabidopsis thaliana* (accession number X15550), *Oryza sativa* (X54194), *Pisum sativum* (X16614), *Vicia sativa* (AY234366), and *Vicia faba* (X16615)], Fungi [*Flammulina velutipes* (MH468771)], Strameopiles [*Fragilariopsis kerguelensis* (LR812489), *Globisporangium ultimum* (AB370108)].

### Annotation

Some of the sequences were partially annotated to indicate the extent of the 3’ETS, 5’ETS, and TR sections within the IGS (*G. gallus*, *H. sapiens*, *P. sativum*, *V. faba*, and *V. sativa*). For this study, additional IGS and ETS TRs ≥20 bp for all species were manually located. Some previously-reported repeats were extended to yield longer head-to-tail TRs. Manual searches for direct repeats (DRs) at both ends of each tandem repeat (TR) section was undertaken. Searches for short DRs (2-10 bp), proposed to be TSDs, were performed manually. For 2 bp DRs, they were counted if they were immediately adjacent to one another (e.g., …ctct…) or separated by a single base pair (e.g., …tgxtg…). For 3 bp DRs, they were counted if they were adjacent or less than 20 bp apart. For longer DRs, they were counted if they were within 40 bp of one another. Potential DRs farther apart than 40 bp were not considered. All of these sections were then mapped (Fig 1) and tabulated (Table 1).

### Statistical Analysis

A program, provided by Luca Comai (University of California, Davis), was used to locate all DRs regardless of the distance from one to another. Analyses of the IGSs and the coding sections were performed separately (Table S1). It also produced a dot plot of all the DRs in the sequence (Fig S1), regardless of the distances between the DRs. It generated a dot plot of 1000 random sequences with the same G+C percentage (Fig S2). It then compared the numbers of DRs in the real sequence versus the random sequences, and then calculated the p-values of whether the number of DRs in the real sequence differed from the number of DRs in each random sequence (Fig S3). The null hypothesis is that they do not differ.

## Glossary

Direct repeats (DRs): repeat sequences (2−10 bp) flanking an unrelated sequence. DRs usually remain as a remnant after a transposon leaves a genomic site.
External transcribed spacer (ETS): the segment following the large rRNA subunit gene (LSU) that is transcribed is the 3’ ETS, whereas the transcribed segment preceding the small rRNA subunit gene (SSU) is the 5’ ETS.
Intergenic spacer (IGS): the spacer segment between the LSU and SSU, also known as the non-transcribed spacer, although it may sometimes be transcribed.
LINEs and SINEs: long (autonomous) and short (nonautonomous) interspersed nuclear elements, respectively (retrotransposons).
MITEs: Miniature inverted-repeat transposable elements.
Repeat Type: R1 is the first type of tandem repeat (TR) sequence in the rDNA spacer following the 3’ETS (see Fig 3). R2 is the second TR type, and so forth. IGSs with only one type of repeat are simply designated R.
Satellite DNA (satDNA): tandem repeats of any unit-length sequence. Here, we designate repeats of <20 bp as DRs and those ≥20 bp as TRs.
Tandem repeats (TRs): typically 10 to >2000 bp that are the defining feature of satellite DNA.
Target site duplication (TSD): duplication of a short sequence at a transposon integration site creating one copy of each that flank the transposon.

## Acknowledgments

We thank George Miklos for his critical comments. The statistical program was written and supplied by Luca Comai.

## Author Contributions

Planning and conceptualization, A.J.B.; Bioinformatic analyses, S.O.R.; Writing and editing manuscript, A.J.B. and S.O.R.; Figures, S.O.R.

## Supporting Information

**Table S1.** Statistical analyses of direct repeats in the intergenic spacers and rRNA genes.

| Species | Region<sup>a</sup> | Stats | Repeat Lengths<sup>b</sup> |  |  |  |  |  |  |  |  |
| --- | --- | --- | --- | --- | --- | --- | --- | --- | --- | --- | --- |
|  |  |  | 2 bp | 3 bp | 4 bp | 5 bp | 6 bp | 7 bp | 8 bp | 9 bp | 10 bp |
| *Oryza sativa* | IGS (2139 bp)<br>72% GC | DR<sup>c</sup> | 202209 | 60571 | 18779 | 6688 | 2996 | 1668 | 1122 | 836 | 697 |
|  |  | p-value<sup>d</sup> | 2.4e-01 | 2.9e-01 | 4.7e-01 | 8.3e-03 | 1.1e-14 | 1.6e-72 | 1.0e-286 | 0.0 | 0.0 |
|  | SSU (358 bp)<br>41% GC | DR | 4188 | 1058 | 225 | 62 | 16 | 4 | 0 | 0 | 0 |
|  |  | p-value | 3.5e-01 | 3.1e-01 | 1.9e-01 | 2.0e-01 | 3.2e-01 | 3.9e-01 | 2.1e-01 | 3.3e-01 | 4.2e-01 |
| *Pisum sativum* | IGS (2981 bp)<br>46% GC | DR | 302708 | 87597 | 27797 | 10634 | 5709 | 4062 | 3372 | 2958 | 2652 |
|  |  | p-value | 7.7e-07 | 1.9e-33 | 1.2e-97 | 0.0 | 0.0 | 0.0 | 0.0 | 0.0 | 0.0 |
|  | SSU (1812 bp)<br>49% GC | DR | 103447 | 26011 | 6534 | 1633 | 395 | 93 | 19 | 5 | 1 |
|  |  | p-value | 4.9e-01 | 4.5e-01 | 4.8e-01 | 4.8e-01 | 2.8e-01 | 2.2e-01 | 1.5e-01 | 3.4e-01 | 3.4e-01 |
| *Arabidopsis thaliana* | IGS (4705 bp)<br>49% GC | DR | 757889 | 216099 | 69357 | 28972 | 16617 | 11958 | 9616 | 7951 | 6654 |
|  |  | p-value | 2.6e-18 | 7.6e-70 | 1.3e-233 | 0.0 | 0.0 | 0.0 | 0.0 | 0.0 | 0.0 |
|  | LSU (485 bp)<br>54% GC | DR | 7392 | 1847 | 435 | 106 | 25 | 3 | 0 | 0 | 0 |
|  |  | p-value | 4.6e-01 | 4.7e-01 | 1.7e-01 | 2.4e-01 | 2.8e-01 | 1.2e-01 | 1.5e-01 | 3.0e-01 | 3.9e-01 |
| *Vicia sativa* | IGS (2976 bp)<br>48% GC | DR | 295617 | 86233 | 28528 | 10974 | 5277 | 3425 | 2714 | 2323 | 2083 |
|  |  | p-value | 7.3e-63 | 2.2e-300 | 0.0 | 0.0 | 0.0 | 0.0 | 0.0 | 0.0 | 0.0 |
|  | SSU/LSU (69 bp)<br>52% GC | DR | 139 | 37 | 9 | 2 | 0 | 0 | 0 | 0 | 0 |
|  |  | p-value | 1.9e-01 | 3.7e-01 | 3.8e-01 | 3.8e-01 | 2.7e-01 | 3.8e-01 | 4.4e-01 | 4.7e-01 | 4.8e-01 |
| *Vicia faba* | IGS (3488 bp)<br>50% GC | DR | 409776 | 114244 | 34193 | 11304 | 4702 | 2651 | 2014 | 1740 | 1591 |
|  |  | p-value | 1.2e-243 | 0.0 | 0.0 | 0.0 | 0.0 | 0.0 | 0.0 | 0.0 | 0.0 |
|  | SSU/LSU (747 bp)<br>GC = 57% | DR | 19019 | 5073 | 1371 | 381 | 103 | 25 | 4 | 0 | 0 |
|  |  | p-value | 4.3e-01 | 2.6e-01 | 1.6e-01 | 9.2e-02 | 1.4e-01 | 3.6e-01 | 2.9e-01 | 1.8e-01 | 3.3e-01 |
| *Flammulina velutipes* | IGS (5198 bp)<br>49% GC | DR | 895662 | 241977 | 69954 | 21664 | 8187 | 4179 | 2930 | 2384 | 2166 |
|  |  | p-value | 9.e-06 | 1.3e-18 | 1.9e-59 | 1.4e-146 | 0.0 | 0.0 | 0.0 | 0.0 | 0.0 |
|  | SSU (1796 bp)<br>46% GC | DR | 103666 | 26383 | 6653 | 1704 | 432 | 109 | 28 | 7 | 0 |
|  |  | p-value | 3.5e-01 | 4.4e-01 | 3.7e-01 | 4.9e-01 | 4.8e-01 | 5.0e-01 | 4.7e-01 | 4.8e-01 | 1.5e-01 |
| *Fragillariopsis kerguelensis* | IGS (4265 bp)<br>33% GC | DR | 726479 | 213321 | 64467 | 19967 | 6580 | 2394 | 1074 | 608 | 439 |
|  |  | p-value | 2.3e-01 | 0.015 | 1.6e-04 | 1.1e-07 | 1.1e-14 | 1.0e-31 | 5.3e-81 | 6.2e-224 | 0.0 |
|  | SSU (744 bp)<br>44% GC | DR | 19034 | 4952 | 1285 | 332 | 85 | 21 | 4 | 2 | 1 |
|  |  | p-value | 3.6e-01 | 4.1e-01 | 4.7e-01 | 4.7e-01 | 4.4e-01 | 3.9e-01 | 2.7e-01 | 4.2e-01 | 2.5e-01 |
| *Globisporangium ultimatum* | IGS (2963 bp)<br>41% GC | DR | 301192 | 84347 | 24234 | 7719 | 2920 | 1460 | 972 | 740 | 610 |
|  |  | p-value | 2.2e-01 | 5.2e-05 | 4.9e-11 | 2.6e-30 | 1.4e-79 | 5.1e-264 | 0.0 | 0.0 | 0.0 |
|  | SSU (1789 bp)<br>45% GC | DR | 104536 | 26872 | 6860 | 1794 | 493 | 132 | 28 | 8 | 1 |
|  |  | p-value | 4.5e-01 | 4.6e-01 | 4.7e-01 | 2.9e-01 | 9.0e-02 | 1.4e-01 | 4.e-01 | 4.7e-01 | 3.0e-01 |
| *Cyanea nozakii* | IGS (1890 bp)<br>49% GC | DR | 119216 | 32209 | 9366 | 3329 | 1697 | 1195 | 993 | 894 | 838 |
|  |  | p-value | 1.5e-02 | 9.0e-06 | 3.5e-16 | 4.5e-64 | 2.9e-286 | 0.0 | 0.0 | 0.0 | 0.0 |
|  | SSU (1815 bp)<br>46% GC | DR | 105518 | 26721 | 6680 | 1643 | 402 | 96 | 19 | 4 | 1 |
|  |  | p-value | 4.5e-01 | 4.4e-01 | 3.8e-01 | 1.9e-01 | 1.7e-01 | 1.9e-01 | 1.1e-01 | 1.9e-01 | 3.2e-01 |
| *Drosophila funebris* | IGS (4810 bp) | DR | 1034905 | 337798 | 118549 | 48261 | 24549 | 17045 | 14400 | 13311 | 12860 |
|  | 30% GC | p-value | 3.3e-02 | 4.0e-08 | 3.5e-24 | 4.2e-80 | 0.0 | 0.0 | 0.0 | 0.0 | 0.0 |
|  | SSU (324 bp) | DR | 3462 | 873 | 240 | 72 | 26 | 8 | 3 | 1 | 0 |
|  | 39% GC | p-value | 2.4e-01 | 1.9e-01 | 4.4e-01 | 2.9e-01 | 6.2e-02 | 1.1e-01 | 1.1e-01 | 1.7e-01 | 4.2e-01 |
| *Gallus gallus* | IGS (17643 bp) | DR | 13288318 | 4254725 | 1454471 | 579847 | 290036 | 183650 | 134846 | 110177 | 95150 |
|  | 70% GC | p-value | 2.3e-01 | 5.0e-08 | 4.5e-37 | 2.8e-163 | 0.0 | 0.0 | 0.0 | 0.0 | 0.0 |
|  | SSU (1823 bp) | DR | 106389 | 27012 | 6905 | 1800 | 446 | 99 | 20 | 3 | 0 |
|  | 54% GC | p-value | 1.9e-01 | 1.6e-01 | 9.4e-02 | 3.9e-02 | 2.4e-01 | 2.7e-01 | 1.4e-01 | 1.3e-01 | 1.6e-01 |
| *Homo sapiens* | IGS (24944 bp) | DR | 23478157 | 7631951 | 2728308 | 1275894 | 673809 | 413327 | 258645 | 180692 | 126720 |
|  | 51% GC | p-value | 2.6e-26 | 4.2e-244 | 0.0 | 0.0 | 0.0 | 0.0 | 0.0 | 0.0 | 0.0 |
|  | SSU (1868 bp) | DR | 113964 | 29235 | 7473 | 1940 | 504 | 133 | 35 | 11 | 3 |
|  | 56% GC | p-value | 1.9e-01 | 1.3e-01 | 1.7e-01 | 1.1e-01 | 1.7e-01 | 2.0e-01 | 2.6e-01 | 1.5e-01 | 2.8e-01 |

<sup>a</sup> IGS = intergenic spacer; SSU = small subunit gene (some are partial sequences); LSU = large subunit gene (partial sequences); SSU/LSU = combination of the SSU and LSU genes (partial sequences).
<sup>b</sup> Direct repeat lengths being compared.
<sup>c</sup> DR = Direct repeats. The number of repeats was determined by creating a matrix with the sequence plotted on the horizontal axis against the same sequence on the vertical axis (see Figs. S1 and S2). Each repeat was then potted against the identical repeat that occurred elsewhere in the sequence. The algorithm then moved to the next nucleotide and performed the same analysis. The total of all such repeats was then recorded, as shown in each column. For example, for *O. sativa* 5-bp repeats, the entire sequence for the IGS is 2139 bp, so the entire matrix created is $2139^2 = 4,575,321$ possible positions (see Fig. S1). A total of 6688 5-bp repeats were identified by the algorithm.
<sup>d</sup> The p-values were calculated by comparing the matrix of repeats based on the actual sequence with 1000 random sequences with the same percent GC (see Figs. S2 and S3). Black font indicates p-values where the actual sequences are significantly different at the $p < 0.01$ level. Red font indicates non-significant differences between the actual sequences and randomly generated sequences. Purple font indicates significant differences at the $p < 0.05$ level. The IGSs all differ from the random sequences at the $p < 0.01$ level, except in *O. sativa*, which exhibits differences from random in direct repeats that are 5 bp and longer. Direct repeat totals in the SSU and LSU sequences did not differ from random sequences at the $p < 0.05$ or $p < 0.01$ levels.

## FIGURES

**Fig. S1.**
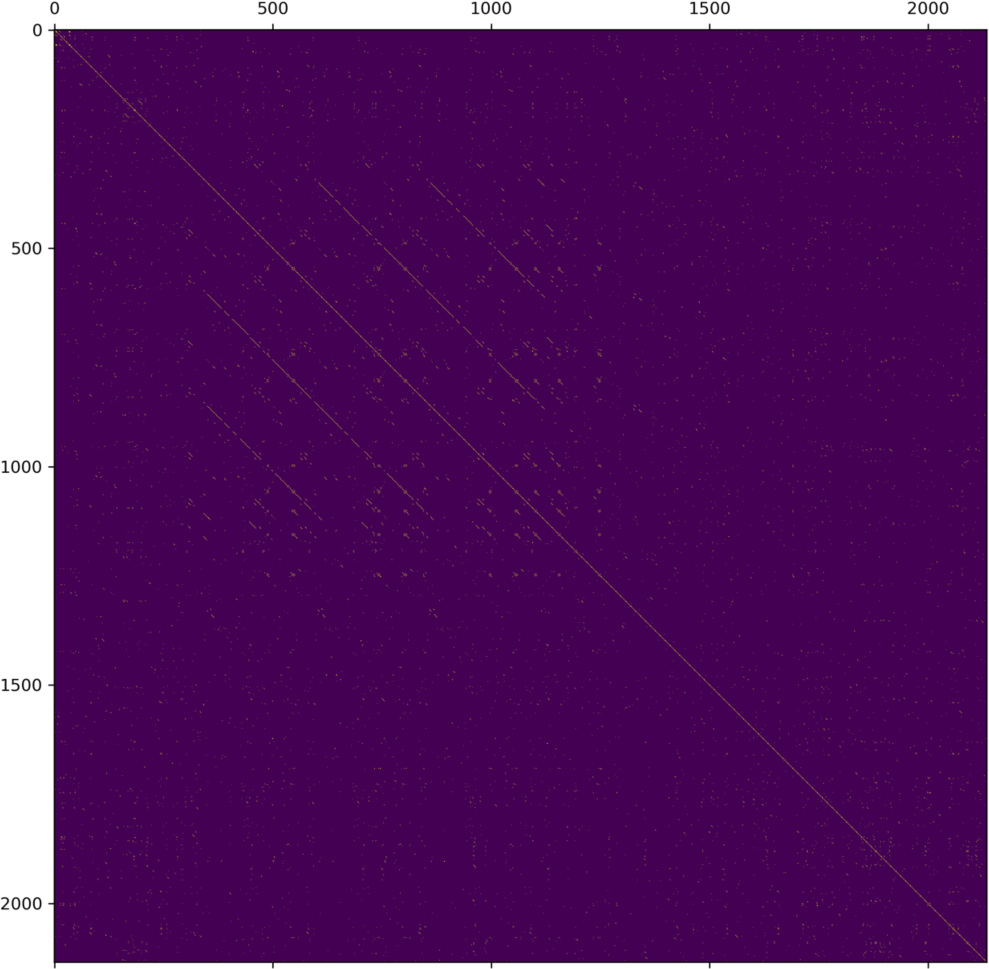
Dot plot of the positions off all 5-bp repeats in the *Oryza sativa* IGS.

**Fig. S2.**
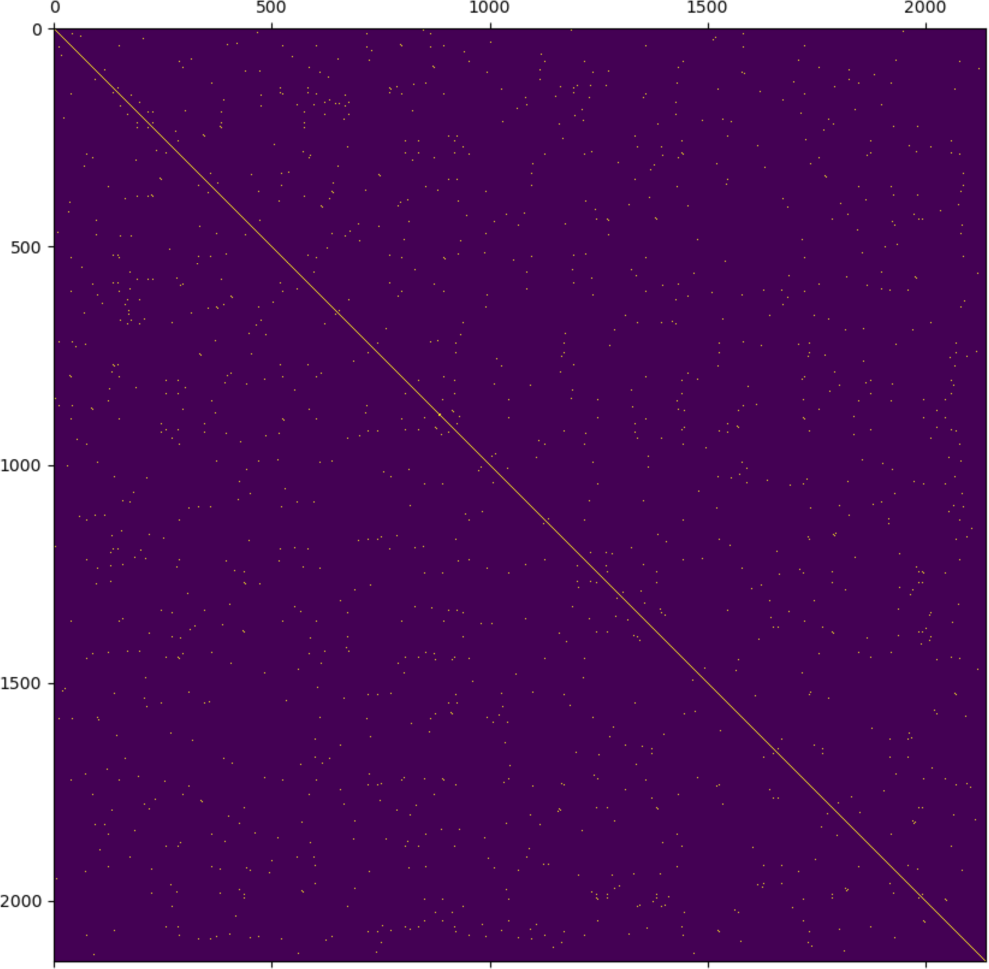
Dot plot of 5-bp repeats in a random sequence.

**Fig. S3.**
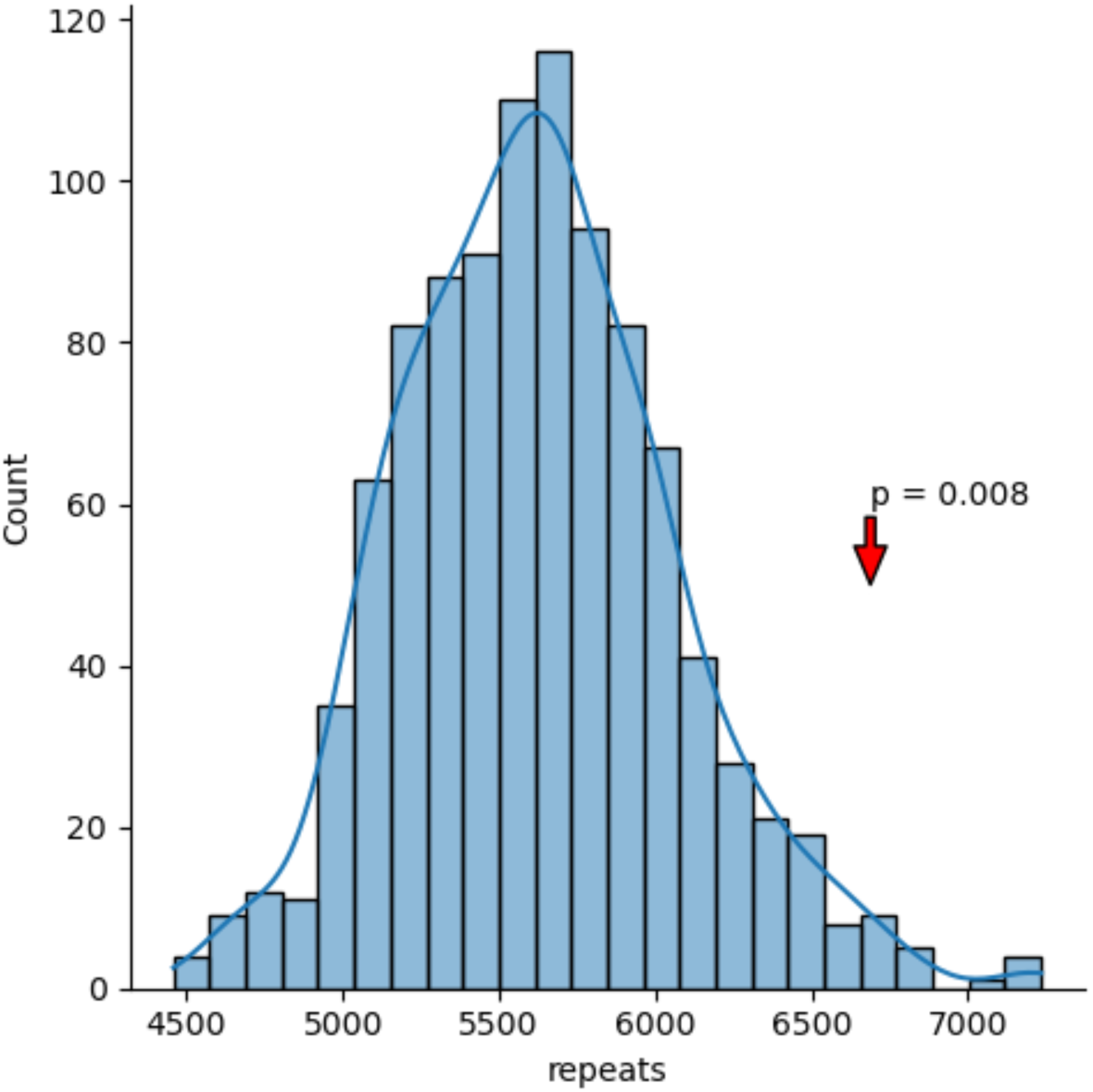
Distribution of 5-bp repeats from 1000 random sequences. Red arrow indicates the actual number of 5-bp repeats in the *Oryza sativa* IGS used in this research.

## References

1. Bendich AJ, McCarthy BJ. Ribosomal RNA homologies among distantly related organisms. Proc. Natl. Acad. Sci. USA. 1970;65:349–356.

2. Pace NR. Mapping the tree of life: progress and prospect. Microbiol. Molec. Biol. Rev. 2009;73:565–576.

3. Rogers SO, Bendich AJ. Ribosomal RNA genes in plants: variability in copy number and in the intergenic spacer. Plant Mol. Biol. 1987;9:509–520.

4. Rogers SO, Bendich AJ. Heritability and variability in ribosomal RNA genes of *Vicia faba*. Genetics 1987;117:285–295.

5. Kasselimi E, Pefane D-E, Taraviras S, Lygerou Z. Ribosomal DNA and the nucleolus at the heart of aging. Trends Biochem. Sci. 2022;47:P328–341. doi.org/10.1016/j.tibs.2021.12.007

6. Lutterman T, Rückert C, Wibberg D, Busche T, Schwarzhans J-P, Friehs K, Kalinowski J. Establishment of a near-contiguous genome sequence of the citric acid producing yeast *Yarrowia lipolytica* DSM 3286 with resolution of rDNA clusters and telomeres. NAR Genom. Bioinform. 2021. doi: 10.1093/nargab/lqab085.

7. Beauparlant MA, Drouin G. Multiple independent insertions of 5S rRNA genes in the spliced-leader gene family of trypanosome species. Curr. Genet. 2014;60:17–24.

8. Dyomin A, Galkina S, Fillon V, Cauet S, Lopez-Roques C, Rodde N, Kloop C, Vignal A, Sokolovskaya A, Satifitdinova A, Gaginskaya E. Structure of the intergenic spacers in chicken ribosomal RNA. Genet. Sel. Evol. 2019;51:59. doi:org/10.1186/s12711-019-0501-7

9. Berger S, Schweiger H-H. Ribosomal DNA in different members of a family of green algae (*Chlorophyta*, Dasycladaceae): and electron microscopical study. Planta 1975;127:49–62

10. Jorgensen RA, Cuellar RE, Thompson WF, Kavanagh TA. Structure and variation in ribosomal RNA genes in pea: characterization of a cloned rDNA repeat and chromosomal rDNA variants. Plant Mol. Biol. 1987;8:3–12.

11. Garrido-Ramos MA. Satellite DNA: an evolving topic. Genes 2017;8:230. doi:10.3390/genes8090230

12. Fort V, Khlifi G, Hussein SM. Long non-coding RNAs and transposable elements: a functional relationship. Mol. Cell Res. 2021;1868:118837 doi.org/10.1016/j.mmamcr.2020.118837

13. Richard G-F, Kerrest A, Dujon B. Comparative genomics and molecular dynamics of DNA repeats in eukaryotes. Microbiol. Mol. Biol. Rev. 2008;72:686–727.

14. Thakur J, Packiaraj J, Henikoff A. Sequence, chromatin and evolution of satellite DNA. Int. J. Mol. Sci. 2021;22:4309. doi.org/10.3390/jms22094309

15. Casacuberta E. *Drosophila*: retrotransposons making up telomeres. Viruses 2017;9:192. doi: 10.3390/v9070192

16. Grandi FC, An W. Non-LTR retrotransposons and microsatellites: partners in genomic variation. Mob. Genet. Elements 2013;3:e25674. doi: 10.4161/mge.25674

17. Tunjić-Cvitanić M, Pasantes JJ, Garcia-Souto D, Cvitanić T, Plohl M, Šatović-Vukšić, E. Satellitome analysis of the pacific oyster *Crassostrea gigas* reveals new pattern of satellite DNA organization, highly scattered around the genome. Int. J. Mol. Sci. 2021;22:6798. doi.org/10.3390/ijms22136798

18. Berger S, Zellmer DM, Kloppstch K, Richter G, Dillard WL, Schweiger HG. Alternating polarity in rRNA genes. Cell Biol. Int. Rep. 1978;2:41–50

19. Franke WW, Scheer, Spring H, Trendelenburg MF, Krohne G. Morphology of transcriptional units of rDNA: evidence for transcription in apparent spacer intercepts and cleavages in the elongating nascent RNA. Exper. Cell Res. 1976;100:233–244.

20. Yakura K, Kato A, Tanifuji S. Length heterogeneity in the large spacer of *Vicia faba* rDNA is due to the differing number of 325 bp repetitive sequence elements. Mol. Gen. Genet. 1984;193:400–405.

21. Kato A, Yakura K, Tanifuji S. Repeated DNA sequences found in the large spacer of *Vicia faba* rDNA. Biochem. Biophys. Acta 1985;825:411–415.

22. Cullis CA, Davies DR. Ribosomal DNA amounts in *Pisum sativum*. Genetics 1975;81:485–492.

23. Ingle J, Timmis J, Sinclair J. The relationship between satellite DNA, ribosomal RNA gene redundancy, and genome size in plants. Plant Physiol. 1975;55:496–501.

24. Pollans NO, Weeden NF, Thompson WF. Distribution, inheritance and lineage relationship of ribosomal DNA spacer length variants in pea. Theor. Appl. Genet. 1986;72:289–295.

25. Havlova K, Dvořáčková M, Peiro R, Abia D, Mozgová I, Vansáčová L, Gutierrez C, Fajkus J. Variation of 45S rDNA intergenic spacers in *Arabidopsis thaliana*. Plant Mol. Biol. 2016;92:457–471.

26. Agrawal S, Ganley ARD. The conservation landscape of the human ribosomal RNA gene Repeats. PLoS ONE 2018;13(12):e0207531. doi:org/10.1371/journal.pone.0207531

27. Meštrović N, Miravinac B, Pavlek M, Vojvoda-Zeljko T, Šatović E, Plohl M. Structural and functional liaisons between transposable elements and satellite DNAs. Chromosome Res. 2015;23:583–596.

28. Rodgers K, McVey, M. Error-prone repair of DNA double-strand breaks. J. Cell Physiol. 2016;21:15–24.

29. Deininger P. *Alu* elements: know the SINEs. Genome Biol. 2011;12:236. doi:org/10.1186/gb-2011-12-12-236

30. Rogers SO. Integrated evolution of ribosomal RNAs, introns, and intron nurseries. Genetica 2019;147:103–119. doi.org/10.1007/s107009-018-0050-y

31. French SL, Osheim, YN, Cioci F, Nomura M, Beyer AL. In exponentially growing *Saccharomyces cerevisiae* cells, rRNA synthesis is determined by the summed RNA polymerase I loading rate rather than the number of active genes. Mol. Cell. Biol. 2003;23:1558–1598.

32. Hori Y, Engel C, Kobayashi T. (2023) Regulation of ribosomal RNA gene copy number, transcription and nucleolus organization in eukaryotes. Nature Rev. Molec. Cell Biol. 2023, doi.org/10.1038/s41580-022-00573-9

33. Salim, D, Bradford WD, Freeland A, Cady G, Wang J, Pruitt S, Gerton J. DNA replication stress restricts DNA copy number. PLoS Genet. 2017;13:e1007006. doi.org/10.1371/journalpgen.1007006

34. Kobayashi T, Horiuchi T, Tongaonkar P, Vu LNomura M. SIR2 regulates recombination between different rDNA repeats, but not recombination within individual rRNA genes in yeast. Cell 2014;117:441–453.

35. Abraham KJ, Khosraviani N, Chan JNY, Gorthi A, Samman A, Zhao DY, et al. Nucleolar RNA polymerase II drives ribosome biogenesis. Nature 2020;585:298–302.

36. Burton RS, Metz EC, Flowers JM, Willett CS. Unusual structure of ribosomal DNA in the copepod *Tigriopus californicus*: intergenic spacer sequences lack internal subrepeats. Gene 2005;344:105– 113. doi:10.1016/j.gene.2004.09.001

37. Koonin EV, Makarova KS, Wolf YI. Evolutionary Genomics of Defense Systems in Archaea and Bacteria. Annu. Rev. Microbiol. 2017;71:233–261.

38. Kirchberger PC, Schmidt M, Ochman H. The Ingenuity of Bacterial Genomes. Annu. Rev. Microbiol. 2020;74:815–834.

39. Gao L, Altae-Tran H, Böjning F, Makarova KS, Segel M, Schmid-Burgk JL, Koob J, Wolf YI, Koonin EV, Zhang F. Diverse enzymatic activities mediate antiviral immunity in prokaryotes. Science 2020;369:1077–1084. doi: 10.1126/science.aba0372

40. Palazzo AF, Gregory TR. The case for junk DNA. PLoS Genetics 2014. doi.org/10.1371/journal.pgen.1004351

41. Condon C, Squires C, Squires CL. Control of rRNA transcription in Escherichia coli. Microb. Rev.1995; 59:623–645.

42. Fleurier S, Dapa T, Tenaillon O, Condon C, Matic I. rRNA operon multiplicity as a bacterial genome stability insurance policy. Nucl. Acids Res. 2022. doi.org/10.1093/nar/gkac332

43. Rogers, S.O. and A.J. Bendich, 1988. Recombination in *E. coli* between cloned ribosomal RNA intergenic spacers from *Vicia faba*: a model for the generation of ribosomal RNA gene heterogeneity in plants. Plant Science 55:27–31.

44. Saini N, Gordenin DA. Hypermutation in single-stranded DNA. DNA Repair (Amst) 2020. doi: 10.1016/jdnarep.2020.102868

45. Pham N, Yan Z, Yu Y, Afreen MF, Malkova A, Haber JE, Ira G. Mechanisms restraining break-induced replication at two-ended DNA double-strand breaks. EMBO .J 2021;e104847. doi.org/10.15252/embj.2020104847

46. Osia B, Twarowski J, Jackson T, Lobachev K, Liu L, Malkova A. Migrating bubble synthesis promotes mutagenesis through lesions in its template. Nucl. Acids Res. 2022;50:6870– 6889. doi.org/10.1093/nar/gkac520

47. de Koning APJ, Gu W, Castoe TA, Batzer MA, Pollack DD. Repetitive elements may comprise over two-thirds of the human genome. PLoS Genet. 2011;7(12):e1002384. doi:10.1371/journal.pgen.1002384

48. Bhowmick R, Lerdrup M, Gadi SA, Rossetti GG, Singh MI, Liu Y, Halazonetis TD, Hickson ID. RAD51 protects human cells from transcription-replication conflicts. Molec, Cell 2022. https://doi.org/10.1016/j.molcel.2022.07.010

49. Bendich AJ, Anderson RS. Characterization of families of repeated DNA sequences from four vascular plants. Biochemistry 1977;16:4655–4663.

50. McClintock, B. (1984) The significance of responses of the genome to challenge. Science 1984;226:792–801.

